# Temperature station matching for elevation-standardised ecological meta-analysis

**DOI:** 10.64898/2026.03.10.709008

**Authors:** Denise Boehnke

## Abstract

1. Standardising temperature data across heterogeneous study sites is essential for ecological meta-analyses, yet elevation-driven lapse rates often confound direct comparisons of coarse-grid climate data. Ecological studies frequently document only site altitude – particularly historical datasets – limiting analysis of thermal influences on spatial organism distribution.
2. A dual-approach protocol was developed to derive regional correction factors (ΔH) from altitude-temperature regressions (Lapse Rate Method: SW Germany/Italian Alps, n=33 stations) and cross-regional station pairs (T^AV^ Matching Method, n=27) with closely aligned long-term mean temperatures (ΔT^AV^ ≤ 1.2°C). Applied to 109 *Ixodes ricinus* study sites across nine European regions, correction factors were calculated only for regions with consistent altitude shifts (ΔH > 100m) relative to Southwest German reference stations.
3. Regional correction factors (ΔH) from both methods included +1300 m (Finland, T^AV^ Matching), +370 m (Netherlands/NE Germany, T^AV^ Matching), and −220 m (Italian Alps, Lapse Rate Method) across five regions. In total, 27 cross-regional T^AV^ matched pairs demonstrated high matching precision (median ΔT^AV^ = 0.05°C, 89 % ≤ 0.2°C). These factors standardised site altitudes to a common SW German thermal reference frame, enabling cross-site comparability.
4. The dual-method protocol requires no automation and is applicable to any taxa with documented site altitudes. The complete methodological workflow - including station data, lapse rate regressions, matching decisions, and correction calculations is publicly available at Zenodo [DOI 10.5281/zenodo.18835116], providing ecologists with a pragmatic, fully reproducible template for elevation-standardised temperature estimation in meta-analyses.

## 1 Introduction

To explore thermal driven effects on population densities at land, data on local temperature are required. These data are often missing or, depending on the research question, available only for the wrong scale, e.g. not representing the organism’s microhabitat [1].

Starting point was a tick density study across SW-Germany in 2014, which functioned as a reference dataset [2,3]. For a meta-analysis investigating the thermal influence on *Ixodes ricinus* density distribution across Europe, 30-year average climate temperature data—or comparable proxies—were required for several study sites. Tick data were drawn from several separated studies across Europa. Those studies only provided the elevation of the tick sampling sites, but no information about long-term temperature.

Large-scale thermal data which provide temperature grids (e.g. E-OBS, European Climate Assessment & Dataset) with the highest resolutions of 0.1° x 0.1° or about 11 km^2^ [4] are unsuitable to represent thermal differences in mountainous terrain.

Thus, a sophisticated methodology was developed to translate the sites’ original altitude into a **site-specific average temperature proxy**.

## 2 Materials and Methods

### 2.1 Population Data

An original tick distribution study with 25 study sites in SW-Germany provided the reference dataset for the meta-analysis [3]. A comprehensive review of European studies on *I. ricinus* resulted in 10 original studies and one data compilation with methodically comparable tick data samples at 84 study sites across Europe (not published yet).

For the meta-analysis, the altitude of all study sampling sites were needed. These were i) drawn from the original papers, ii) refer to a personal request [5] or iii) were calculated by coordinates using a web-based calculation under https://www.gpskoordinaten.de/ for [6,7].

### 2.2 Altitude-temperature relation for fixed latitude

The temperature lapse rate—the decrease in air temperature with increasing altitude at constant latitude—typically ranges from −0.54 to −0.58°C per 100 m and exhibits seasonal variation, with steeper gradients during summer months [8]. This empirically determined relationship form the foundation for using altitude as a temperature proxy within physiographically similar regions.

The naturally strong correlation between altitude and average temperature, also known as *altitudinal temperature gradient* [9] was analyzed via linear regression using long-term temperature averages (1981 to 2010) from a total of 34 official weather stations located in SW Germany (Black forest mountain range) and in the Italian Alps - nearby the study sites of an Italian study that sampled ticks at a broad altitudinal range [5]. Raw station data, matching workflow, and calculated correction factors are available at Zenodo [https://zenodo.org/records/18835117].

### 2.3 Latitude-temperature relation for regional standardisation

Using altitude as a proxy for average temperatures across European regions has an important peculiarity: at the same altitude, T_AV_ is much warmer in the south and much cooler in the North. Or in other words, as shown in the results section Figure 3, the T_AV_ – altitude relationship shifts to higher altitudes in the south compared to the north. Due to this *latitudinal temperature gradient*, altitude can serve as a reliable temperature-proxy only within a limited regional frame.

To derive correction factors accounting for latitudinal temperature gradients, locations with comparable long-term mean temperatures (T_AV_) were identified from official meteorological records within the nine study regions across Europe. For discrete T_AV_ intervals (3.5–4°C; 8.3–8.7°C; 9.8–10.2°C), the corresponding altitudes were determined and plotted against latitude to visualize systematic altitude shifts between regions (Figure 3).

This analysis confirmed that altitude-temperature relationships are regionally consistent but vertically offset by latitude, providing the basis for subsequent station matching and correction factor derivation.

### 2.4 Dual-approach temperature standardisation

The workflow for regional temperature standardisation is illustrated in Figure 1. To account for regional differences in long-term mean temperature (T_AV_) at comparable altitudes, additional official weather stations were compiled for each of the nine study regions. For each target region, stations located near the tick study sites were identified based on geographic proximity. Two complementary approaches - the Lapse Rate Method and the T_AV_ Matching Method - were applied depending on regional station density and physiography.

**Figure 1.**
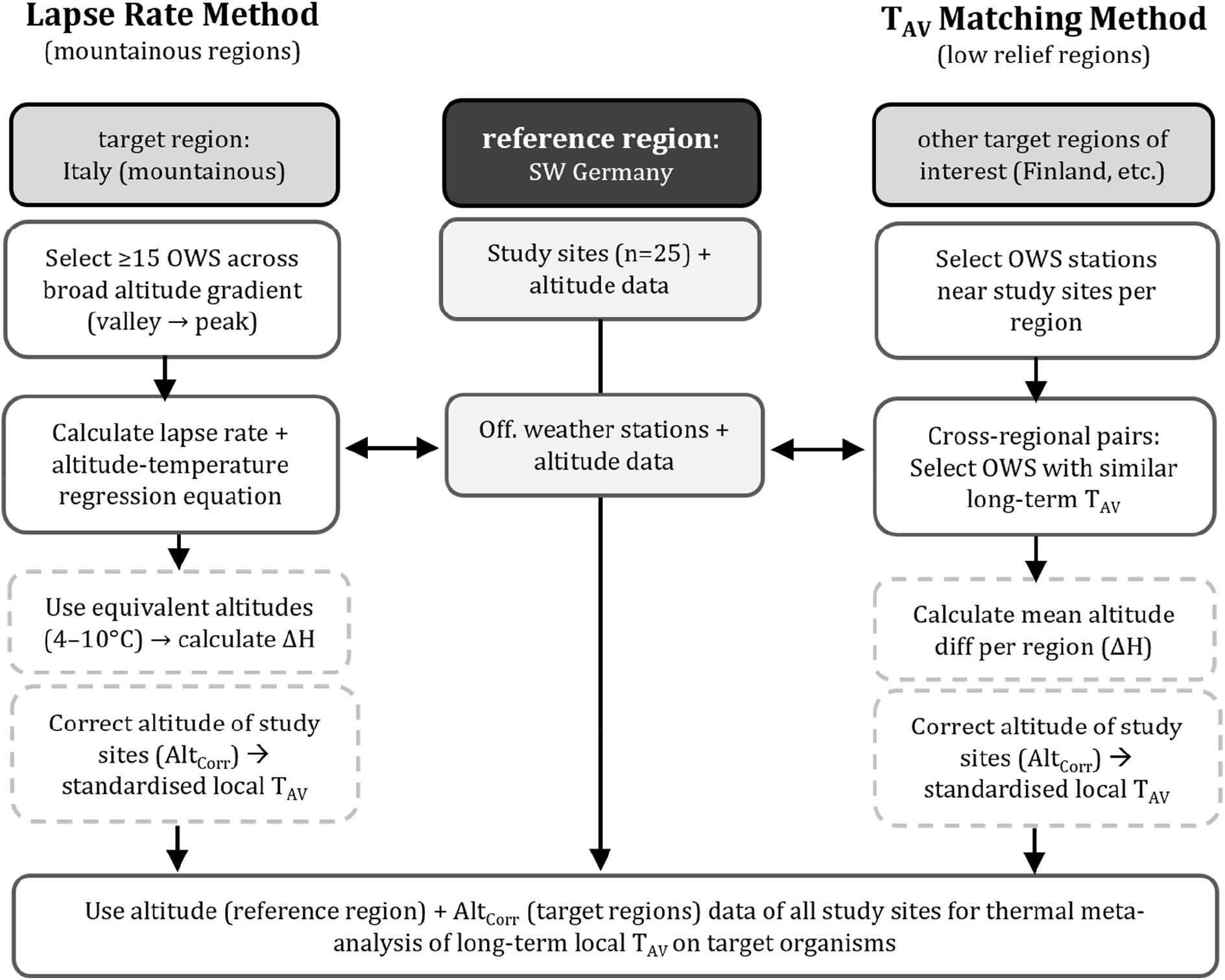
Dual-approach temperature standardisation workflow. Reference region (middle): SW Germany. Lapse Rate Method (Left) especially for mountainous target regions with dense station network (n≥15); long-term temperature (T_AV_) Matching Method (right) for low relief target regions or limited number of stations. OWS = Official Weather Station, diff = difference, ΔH = mean altitude offset.

#### Lapse Rate Method (mountainous regions)

For regions with sufficient station coverage across elevation gradients (SW Germany Black Forest/Swabian Alps: n=18 stations; Italian Alps: n=15 stations), separate altitude- temperature regressions were calculated (valley-to-peak). Equivalent altitudes for discrete T_AV_ intervals (4°C, 5°C,..10°C) were derived from both regression equations, yielding 7 altitude pairs. Regional correction factors (ΔH) represent mean altitude differences between regression lines at identical T_AV_ values. The T_AV_ intervals (4-10°C) reflect the long-term temperature range of the reference region SW Germany.

#### Temperature (T_AV_) Matching Method (low-relief/sparse data regions)

For target regions with few study sites or flat terrain (Finland, Netherlands, NE Germany, Swiss Wallis), target stations were matched directly to SW German reference stations by minimising ΔT_AV_. Station pairs were accepted if ΔT_AV_ ≤ 1.2 °C. Regional correction factors (ΔH > 100 m) were calculated as mean altitude differences across accepted station pairs (n=27). Correction factors were derived only for regions showing consistent altitude shifts (ΔH > 100 m) at comparable T_AV_. Regions with inconsistent or negligible altitude differences were considered climatically aligned with the reference region.

#### Correction factor application

Applying regional ΔH values derived from both approaches - the Lapse Rate Method and the T_AV_ Matching Method - to study sites in all corrected regions produces elevation-standardised altitudes (**Alt**^**corr**^) as **site-specific temperature proxies** within the SW German reference frame, enabling thermal meta-analysis.

See Zenodo files “6-Italy_Germany” (Lapse Rate Method) and “7-Temp_Europe” (T_AV_ Matching Method + application) [https://zenodo.org/records/18835117]. Note: Tick study site locations and Altcorr calculations form the basis for a subsequent *Ixodes ricinus* meta-analysis publication.

## 3 Results

The altitude and long-term mean temperature (T_AV_) relationships confirmed altitude as a reliable temperature proxy within physiographically similar regions.

### 3.1 Altitude-temperature relation and **L**apse **R**ate **M**ethod

To verify lapse rate robustness and derive regional correction factors (ΔH), altitude-T_AV_ relationships were analysed for mountainous regions. In both SW Germany (Black Forest/Swabian Alps, n=18 stations) and the Italian Alps (n=15 stations), T_AV_ declined linearly with altitude, showing nearly identical lapse rates but a systematic vertical offset between regions (Figure 2). Italian Conditions were consistently warmer, corresponding to an altitude offset ΔH=−220 m relative to SW Germany at comparable T_AV_.

**Figure 2.**
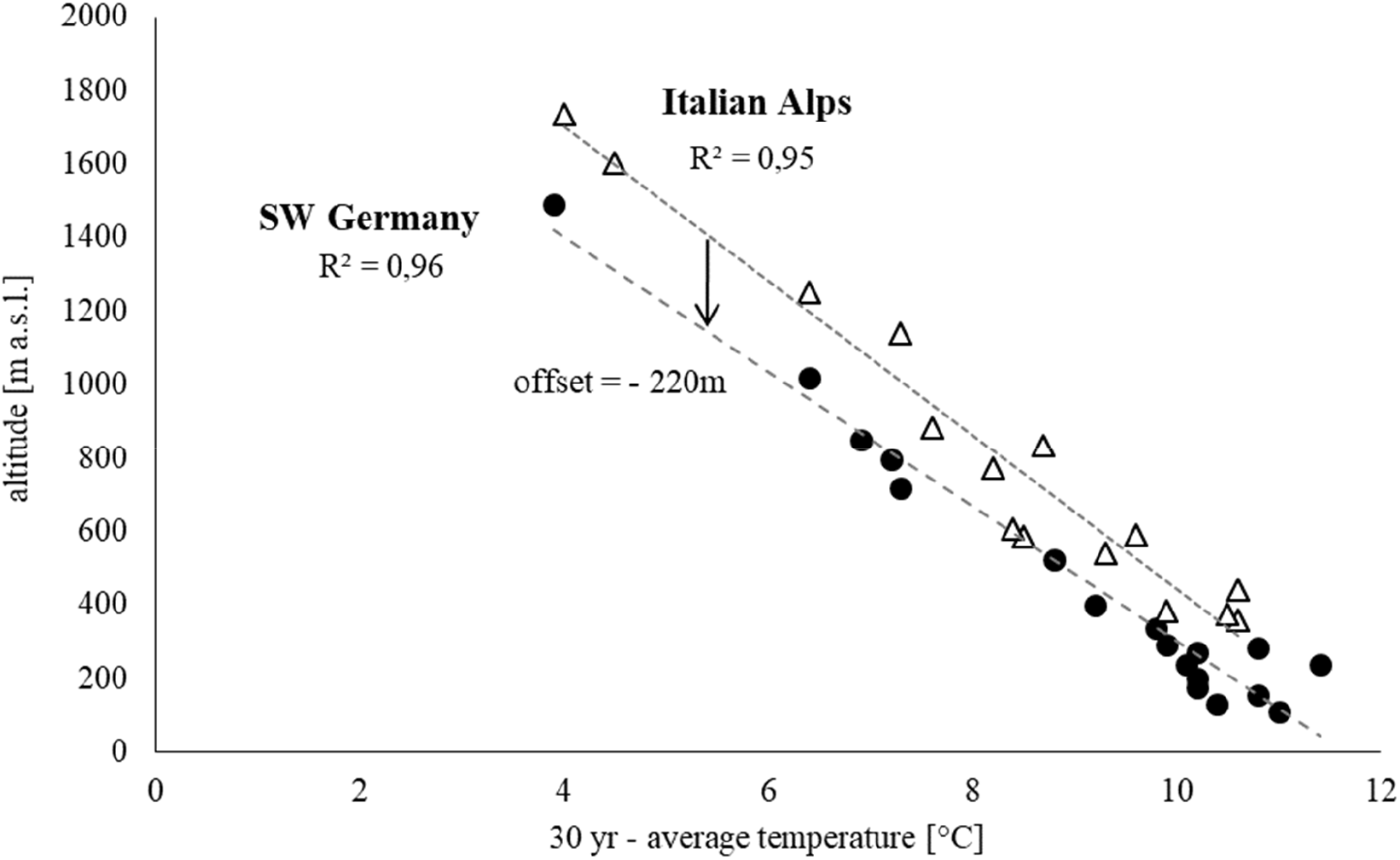
The strong linear relationship of long-term average temperatures (T_AV_) and altitude. Analysis are based on data from official meteorological stations in the Black Forest in SW-Germany and in the Italian Alps. It is also visible that conditions are warmer in Italy, leading to an altitude-offset [ΔH] of about 220m between the two regions.

The strong correlation between altitude and 30yr-average temperatures (T_AV_) together with the parallel regression slopes despite regional offset, underline the validity and robustness of the Lapse Rate Method. This approach is particularly robust when stations span complete elevation gradients from valley bottoms to mountain peaks, enabling derivation of representative lapse rates.

### 3.2 Latitude-temperature relation and T_AV_ matching method

To verify temperature matching robustness and derive cross-regional correction factors (ΔH), T_AV_-equivalent station pairs were identified across Europe. Sites with comparable T_AV_ occurred systematically at higher altitudes in southern regions than in northern regions. Along the European latitudinal gradient, equivalent T_AV_ values shifted by approximately +1300 m between SW Germany and Finland and +1700 m relative to the Italian Alps (Figure 3). Regression slopes for discrete T_AV_ intervals remained nearly identical across latitudes, confirming the transferability of regional lapse rates despite systematic altitude offsets.

Regional altitude shifts were visualised by plotting altitude-T_AV_ relationships for the reference region (SW Germany), Italian reference stations, the accepted station pairs (ΔT_AV_ ≤ 1.2°C), and all additional weather stations within each target region (Figure 4).

**Figure 3.**
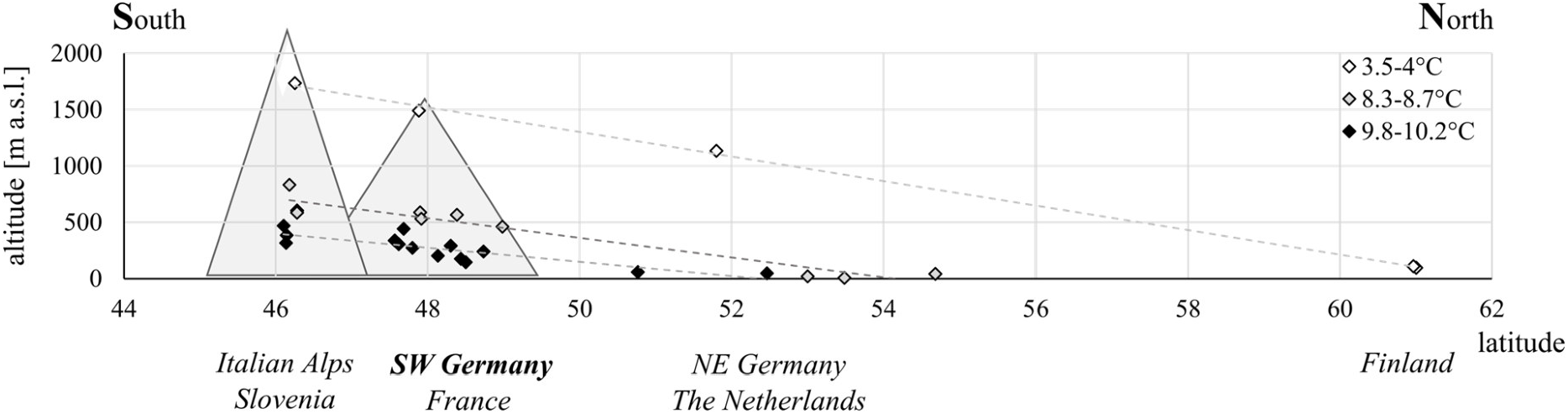
The linear trend of the latitudinal temperature gradient across Europe. Locations with comparable long-term average temperatures (T_AV_, shown as rhombuses) are interlinked by dashed lines, showing a linear trend of temperature decrease.

**Figure 4.**
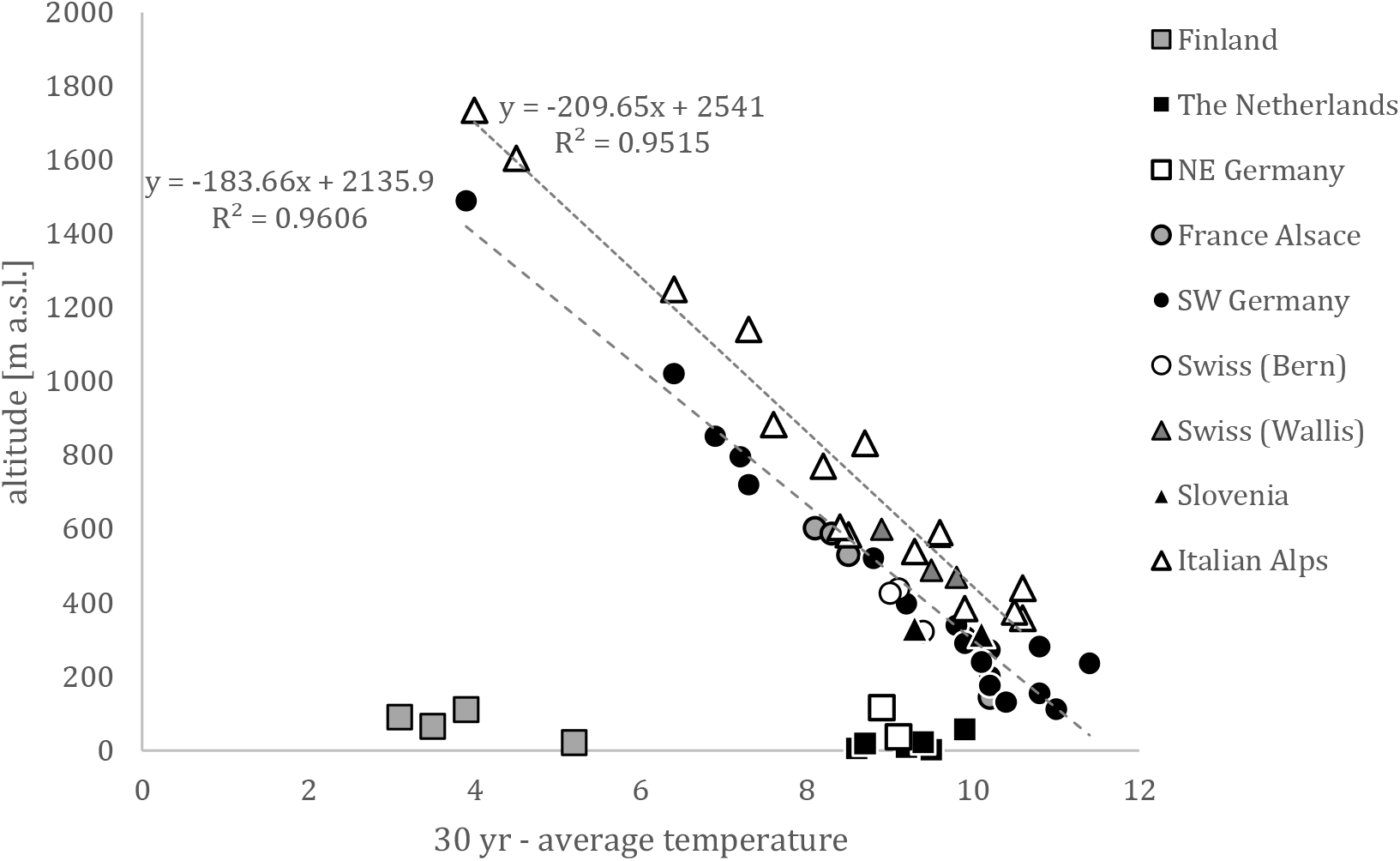
Altitude-T_AV_ relationships across European regions. Reference stations from SW Germany and selected weather stations per target region (compare legend). Consistent latitudinal altitude shifts are evident across the full station network.

Figure 4 validated the consistency of altitude-temperature relationships across regions, and that stations systematically deviating from the SW Germany reference regression line (black dots) required correction, while those proximal to the reference line were climatically equivalent consistent with prior calculations.

### 3.3 temperature standardisation

Regional correction factors (ΔH) were derived from both methods across nine study regions, with consistent altitude shifts (>100 m) identified in five regions (Table 1). Regional ΔH values were rounded to the nearest 10 m to account for methodological uncertainty and prevent over-correction.

**Table 1.** Regional correction factors (ΔH) for temperature standardization, derived from from both methods relative to SW Germany.

| Target region | $\Delta H$ (m) | n pairs | Example shift |
| --- | --- | --- | --- |
| Finland | + 1300 | 3 | 111 --> 1411 m |
| The Netherlands | + 370 | 7 | 25 --> 395 m |
| NE Germany | + 370 | 2 | 117 --> 487 m |
| Swiss (Wallis) | - 90m | 4 | 489 --> 399 m |
| Italian Alps | -220m | 7 | 770 --> 550 m |
| France Alsace | - | 5 | Inconsistent |
| Swiss (bern) | - | 3 | Inconsistent |
| Slovenia | - | 3 | Inconsistent |

#### Lapse Rate Method (Italian Alps)

Altitude equivalents for discrete T_AV_ intervals (4–10°C) between SW Germany and Italian Alps yielded ΔH = −223 m (mean across 7 pairs; Table 2). This systematic offset reflects warmer Italian conditions at equivalent altitudes.

**Table 2.** Italian Alps correction factor (ΔH, Lapse Rate Method) based on mean altitude differences (DE=Germany, IT=Italy) at identical T_AV_.

| $T_{AV}$ (°C) | H_DE (m a.s.l.) | H_IT (m a.s.l.) | $\Delta H$ (m) |
| --- | --- | --- | --- |
| 10 | 299.3 | 444.5 | -145.2 |
| 9 | 482.96 | 654.15 | -171.19 |
| 8 | 666.62 | 863.8 | -197.18 |
| 7 | 850.28 | 1073.45 | -223.17 |
| 6 | 1033.94 | 1283.1 | -249.16 |
| 5 | 1217.6 | 1492.75 | -275.15 |
| 4 | 1401.26 | 1702.4 | -301.14 |
| Mean |  |  | -223.17 |

#### T_AV_ Matching Method (remaining regions)

In total, 27 cross-regional station pairs met the selection criterion of ΔT_AV_ ≤ 1.2°C, yielding ΔH for four additional regions (Finland, Netherlands, NE Germany, Swiss Wallis; Table 1). These regional ΔH values define the equivalent altitude in the reference region for a given target-region. Three regions (France Alsace, Swiss Bern, Slovenia) were excluded due to inconsistent or negligible altitude offsets (<100 m).

The distribution of ΔT_AV_ for the 27 matched station pairs was strongly skewed towards very small mismatches, with a median of 0.05°C and a maximum of 1.2°C (Figure 5). Overall, 89% of pairs showed ΔT_AV_ ≤ 0.2°C, confirming the effectiveness of the minimisation approach for cross-regional temperature matching.

**Figure 5.**
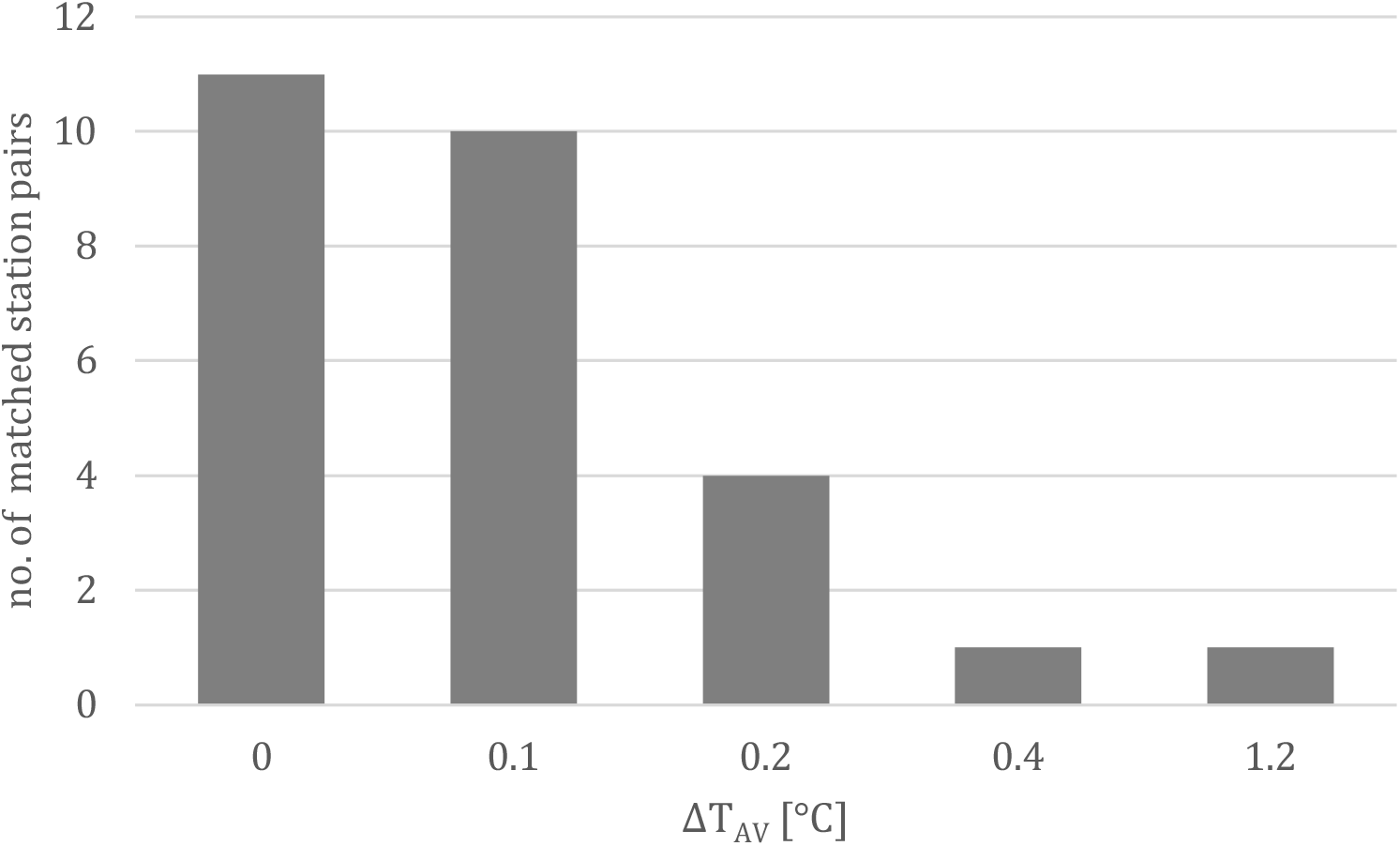
Distribution of ΔT_AV_ for matched station pairs (n=27).

#### Standardisation outcome

Regional ΔH values, derived from both methods, were applied to tick study sites in the five corrected regions, producing elevation-standardised altitudes (Altcorr) within the SW German reference frame. Study sites from colder northern regions were shifted to lower equivalent altitudes, while sites from warmer southern regions were shifted upwards. These corrected values provide a consistent reference frame for thermal meta-analysis across all tick study sites.

### 3.4 Exemplary application: cross-European thermal normalisation

An illustrative application of the standardisation protocol is shown for three *Ixodes ricinus* datasets from Southwest Germany, Finland, and the Italian Alps (Figure 6).

**Figure 6.**
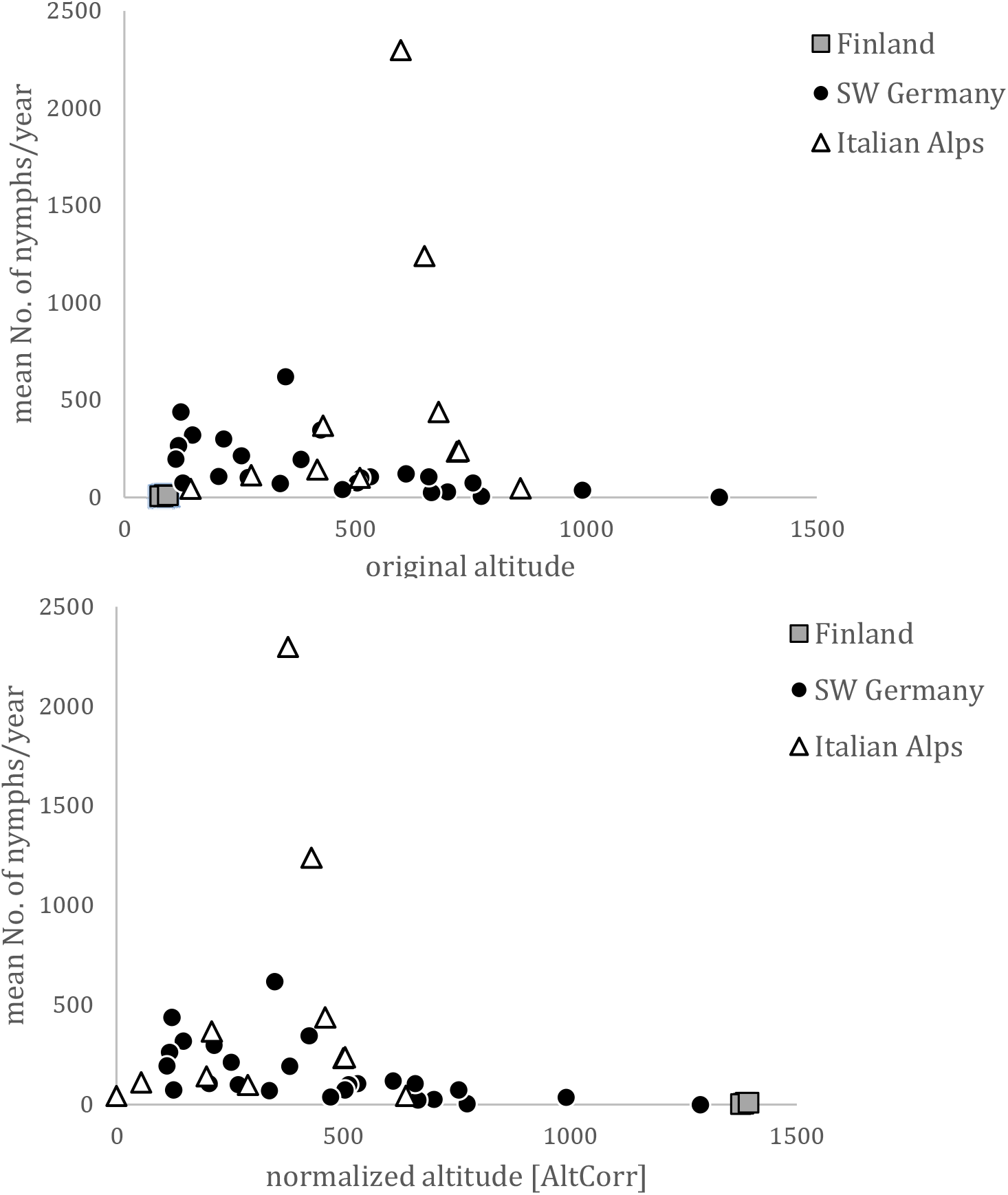
Application of the temperature standardisation protocol to *Ixodes ricinus* nymph density data from Finland (squares), Southwest Germany (circles), and the Italian Alps (triangles). Top panel: Nymphal density plotted against original altitude, showing region-specific elevation ranges. Bottom panel: Altcorr (SW Germany reference frame) after regional ΔH correction, enabling cross-European thermal comparison.

In the original data, nymph densities decline with increasing altitude within regions as expected, but between-region patterns are difficult to compare because sites from colder northern (Finland) and warmer southern regions (Italian Alps) occupy different absolute elevation ranges. After applying regional correction factors (ΔH) and converting site altitudes to the Southwest German reference frame (Altcorr), the three datasets align along a common thermal altitude gradient. Data are provided in the Supplemental Table 1.

This cross-European normalisation enables quantitative analyses of nymph density responses to equivalent thermal conditions across climatically diverse regions, demonstrating the protocol’s utility for advanced meta-analytical applications.

## 4 Discussion

The dual-method protocol (Lapse Rate Method + T_AV_ Matching Method) provides a transparent, pragmatic approach for deriving elevation-standardised temperature estimates from study sites where only altitude data are available.

By deriving regional correction factors from parallel altitude-T_AV_ regressions (ΔH = −220 m, Italian Alps), the approach captures systematic vertical offsets in mountainous terrain with high precision (Lapse Rate Method, Table 2). In addition, minimising ΔT_AV_ between reference and target stations (median 0.05°C, 89% ≤ 0.2°C across n=27 pairs) achieved precise cross-regional alignment in low-relief/sparse-data regions without complex geospatial interpolation (T_AV_ Matching Method, Figure 5).

The selective application of correction factors (>100 m threshold) across both methods prevents overcorrection in climatically similar regions, ensuring methodological robustness. Three regions were excluded due to inconsistent/insufficient offsets. Regional lapse rate consistency across diverse physiographic zones (Black Forest/Swabian Alps, Italian Alps) validates altitude-temperature transferability, while T_AV_ Matching extends this principle to lowlands – creating a comprehensive standardisation framework for Europe’s topographic diversity.

While fully automated matching would be feasible with programming tools, manual selection ensured full traceability of decisions while building methodological confidence and experiential insight into regional differences. The resulting corrected altitudes (Alt^corr^) values enable direct thermal comparability across all 109 study sites and regions, facilitating elevation-stratified meta-analysis of ecological data.

Note that the empirically determined relationships between temperature, altitude, and latitude represent coarse approximations of physical reality, neglecting regional effects such as Gulf Stream influences or local topography. Yet, the protocol successfully normalised diverse datasets into a consistent European reference frame, allowing for European-wide *Ixodes ricinus* meta-analysis.

The protocol is particularly valuable for mountainous regions where coarse-grid satellite data (e.g. E-OBS 0.1°) fail to capture lapse rate variability along elevation gradients. While regional station density remains a prerequisite, the protocol requires minimal computational resources and leverages existing meteorological archives.

Applicable beyond ticks to any taxa with documented site altitudes, it offers ecologists a straightforward template for harmonising heterogeneous climate data—especially in topographically complex landscapes. Meteorological data, calculations, and validation graphics are provided at Zenodo [https://zenodo.org/records/18835117] to support replication and extension to other study systems.

## Data availability

The comprehensive Excel workbook documenting all station data, preselection, matching protocol, latitude/altitude correction factors, and regression graphics is archived at Zenodo [DOI <u>10.5281/zenodo.18835116</u>].

## Acknowledgments

I am very grateful to Stefan Hinz for helpful discussions and valuable suggestions during the development of this station matching protocol. I also acknowledge the valuable contributions of Perplexity AI (powered by Grok 4.1, March 2026 version) for assistance in refining the manuscript structure, text formulation, and logical consistency during final revisions. Station data compilation supported by national meteorological services (DWD and other public weather data providers).

